# Transcriptomic response to host and non-host plants during oviposition in two closely related moth species

**DOI:** 10.1101/244806

**Authors:** M. Orsucci, P. Audiot, S. Nidelet, F. Dorkeld, A. Pommier, M. Vabre, D. Severac, M. Rohmer, B. Gschloessl, R Streiff

**Affiliations:** CBGP UMR 1062, INRA-IRD-CIRAD-Montpellier SupAgro, Montferrier sur Lez, France; DGIMI UMR 1333, INRA-Université de Montpellier, Montpellier, France; MELGUEIL DIASCOPE UE 0398, INRA, Mauguio, France.; present address: Department of Ecology and Genetics, EBC, Uppsala University, Norbyvägen 18D, 75236 Uppsala, Sweden.; MGX-Montpellier GenomiX, c/o Institut de Génomique Fonctionnelle, 34094 Montpellier Cedex 5, France.

## Abstract

We present here a comprehensive analysis of the transcriptomic response to plant environments in ovipositing females of two sibling species of phytophagous moths affiliated to different host ranges: the European corn borer (ECB) and the adzuki bean borer (ABB). We first assembled and annotated a *de novo* reference transcriptome based on a high throughput RNA sequencing of females placed in different plant environments, then we measured differences in gene expression between ECB and ABB, and also within each moth species between environments. We further related the differentially expressed (DE) genes to the host preference in ECB and ABB and highlighted the functional categories involved. More specifically, we conducted an analysis on chemosensory genes previously characterized in ECB, ABB and other related *Ostrinia* species, as these genes are considered as good candidates for the host recognition before oviposition.

Overall, we recorded more DE genes in ECB than in ABB samples, what could highlight the higher strength of the host specialization in ECB compared to ABB as observed at the behavioral level. We also noticed that the genes involved in the preference for their respective host were different between ECB and ABB. At the functional level, the response to plant environment in ECB and ABB during oviposition involved many processes, including the chemosensory repertoire as expected, but also metabolism of carbohydrates, lipids, proteins, and amino acids, detoxification mechanisms and immunity.

All together, our results allowed identifying genes and functions candidates for specialization and also for the species divergence between ECB and ABB. By ad-hoc categorization, we discriminated some genes responding to the environment with similar or divergent pattern in ECB and ABB. Among them, we highlighted new lines of research like carbohydrates metabolism or virus and retrovirus dynamics.

## Introduction

The speciation process by which populations diverge and become reproductive isolated may be driven by divergent selection between different environments that affects various life history traits linked directly or indirectly to fitness (Coyne and Orr, 2004; Funk, 1998; Gavrilets, 2004; Nosil *et al*., 2007; Schluter, 2000; Schluter, 2001). In phytophagous insects, host races are relevant examples of population divergence driven or reinforced by host plant specialization that may, in some cases, be a key component of reproductive isolation (Drès and Mallet, 2002; Forister *et al*., 2012; Futuyma and Moreno, 1988; Matsubayashi *et al*., 2010a). Host plants affect directly the fitness of herbivores (Awmack and Leather, 2006) due to their role in nutrition (Drès and Mallet, 2002; Scriber and Slansky, 1981), protection (Gross, 1993; Larsson *et al*., 1997), or reproduction (Awmack and Leather, 2006; Wood and Keese, 1990). Indeed, plant metabolites are known to affect the partners encounter and mating success (Dupuy *et al*., 2017; Pregitzer *et al*., 2012), the level of egg laying (Foster and Howard, 1999; Leather *et al*., 1985), the choice of oviposition site of gravid females (Renwick, 1994), but also the development of eggs and larvae (Moreau *et al*., 2006). In addition to intrinsic host plant qualities, associated higher-trophic interaction levels such as plant-associated microorganisms and parasitoids may also impact the realized fecundity by reducing the larval performance (Babikova *et al*., 2014). The herbivorous host range is the result of these complex pluritrophic interactions occurring throughout the entire life cycle of the insects (Moreau *et al*., 2006).

In holometabolous species, such as Lepidoptera, the oviposition choice is crucial because, before the imago stage, individuals have often a low mobility and thus depend on the judicious choice of plant by the adult female (Renwick, 1994). The preference–performance hypothesis (also called “mother-knows-the-best” hypothesis) suggests that host preference of adult females should maximize the fitness of their offspring by ovipositing on the most suitable hosts (Jaenike, 1990). Empirical data do sustain this hypothesis for a majority of the study cases (Gripenberg *et al*., 2010) and molecular and functional studies highlighted the importance of chemosensory mechanisms for the recognition by ovipositing females of olfactory cues emitted by the plant during the egg laying step (Carrasco *et al*., 2015; Ma *et al*., 2016). Interestingly, Trona *et al*. (2013) underscored that, in nature, sex pheromone and plant odors are perceived and coded as an ensemble, and suggest a dual role of plant signals in habitat selection and in premating sexual communication.

The European corn borer (ECB, *Ostrinia nubilalis*) and the adzuki bean borer (ABB, *Ostrinia scapulalis*), *sensu* Frolov *et al*. (2007), are two sibling species of phytophagous moths, co-occurring in large part of the European continent. These two species are specialized to different host ranges: ECB is mainly observed on an agricultural monocotyledon, the maize (*Zea mays*) and ABB on several dicotyledons like mugwort (*Artemisia vulgaris*), hop (*Humulus lupulus*) and hemp (*Cannabis sativa*; Frolov *et al*. 2007). In ECB and ABB, the preference-performance hypothesis is typically observed according to various sampling and independent studies (Calcagno *et al*., 2007; Malausa *et al*., 2008; Orsucci *et al*., 2016), but both species exhibit contrasted behaviors and strategies. In a previous experiment, ECB and ABB adult moths were exposed to three plant environments during their mating and egg laying phases: a pure maize environment, a pure mugwort environment and a mixed environment with maize and mugwort plants (Orsucci *et al*., 2016). The authors observed a strong preference for maize in ECB gravid females when they had the choice between maize and mugwort, coupled with a significant avoidance for mugwort plants when this plant was the only possible choice (see Box 1 summary for details). In contrast, ABB showed a moderate preference for mugwort over maize during oviposition and no avoidance behavior.

#### Box 1 Experimental framework of RNAseq sequencing: material, methods and main results on the behavior and life history traits of ECB and ABB during the oviposition phase (inspired by Orsucci *et al* 2016)

#### 1. Experimental design

ECB and ABB larvae were collected near of Versailles, France (48°48019″N, 2°08006″E) from maize and mugwort stands, respectively (Fig B1.A). Mating and growth of the next generations were conducted in laboratory conditions according to (Orsucci *et al*, 2016). After the fifth instar molting, each F2 pupae was isolated until emergence to ensure adult virginity before use in experiments (Fig B1.B). Twenty females and 15 males were released in three different experimental conditions: ‘maize’ (*i.e*. pure maize), ‘mugwort’ (*i.e*. pure mugwort), and ‘choice’ (*i.e*. mixture of mugwort and maize) for the oviposition experiment. After 3 nights (~72h) in semi-natural conditions, behavioral and phenotypic traits were measured. Then, the recovered adults were flash-frozen in nitrogen liquid and stored at -80°C (Fig B1.C). Finally, we realized the dissections (each frozen female was cut in two parts to separate head and thorax tissues from abdomen).

One equimolar pool of RNA extracts was done per cage (Fig B1.D). 24 RNA libraries corresponding to 2 moth species x 3 experimental conditions x 2 repetitions were sequenced in a 1x50bp design on a HiSeq2000 (Illumina).

**Figure B1.**
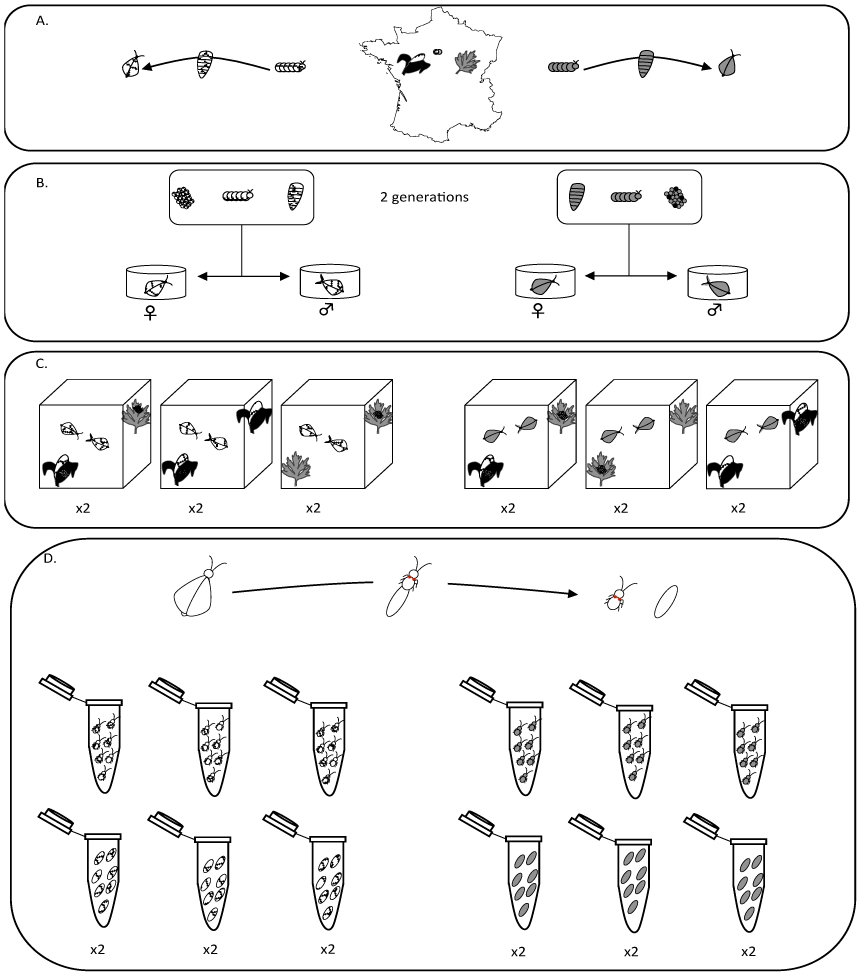
- Experimental design from sampling in natural populations to RNA extractions. ECB and ABB are represented in white and gray, respectively.

#### 2. Summary of life history traits variations in ECB and ABB females during, oviposition

We measured some life history traits variations in ECB and ABB gravid females in the three experiment conditions: ‘maize’, ‘mugwort’, and ‘choice’:

i. *the fertility* was estimated as the total number of eggs laid in cages;
ii. *the host preference for oviposition* was significant when ECB laid more eggs on maize than on mugwort plants in choice conditions, and inversely for ABB;
iii. *the host avoidance for oviposition* was significant when ECB (ABB) females laid more eggs on neutral support (cage netting) than on plants when they had no choice and only mugwort (maize) plants available;
iv. *the survival*, was measured as the number of moths recovered after 3 days of release in experimental cages over the number of moths initially released;
v. *Resting site* concerned the site of capture: mugwort, maize or cage netting.

The results observed for these traits are summarized in Table Bl.

**Table B1.** Main results of oviposition experiment: +/- signs (and YES/NO) indicate the direction and the strength of significant relationships between traits, moth species and experimental conditions. The ‘=’ sign indicates no effect. (Inspired from Orsucci *et al*. 2016).

| TRAIT | ECB |  | ABB |  |
| --- | --- | --- | --- | --- |
|  | MAIZE | MUGWORT | MAIZE | MUGWORT |
| FERTILITY | ++ | -- | - | + |
| HOST PREFERENCE FOR OVIPOSITION | +++ | --- | - | + |
| HOST AVOIDANCE FOR OVIPOSITION | NO | YES | NO |  |
| SURVIVAL | + | - | = |  |
| RESTING | + | - | = |  |

In addition to the oviposition behavior, a higher survival and egg-laying rate in ECB was observed when maize was in its environment, with a preference for maize plants also as resting site. In ABB, these latter traits were not affected by the plant environment (see Box 1 summary). These results suggested the occurrence of attractive and/or repellents molecules impacting the oviposition choice (Leppik and Frérot, 2012) as well as synergetic effects on the mating and egg laying success (Landolt and Phillips, 1997). The olfactory responses of ECB females for plant odors have been previously characterized when presented to: *i)* uninjured and injured maize plants (Schurr and Holdaway, 1970), *ii)* blends of maize volatiles in specific ratios (Molnár *et al*., 2015; Solé *et al*., 2010), *iii)* maize *vs*. mugwort or hop odors (Leppik and Frérot, 2012). However, the underlying molecular mechanisms (*e.g*. genes) of the plant recognition and choice are still unknown.

We present here a comprehensive gene expression analysis of ECB and ABB adult females through an high throughput RNA sequencing of females placed in various plant environments during the oviposition step (sampled from the experiment in Orsucci *et al*., 2016, see Box 1 experimental design for details). We assembled and annotated a *de novo* reference transcriptome and measured differences in gene expression, first between ECB and ABB, then within each moth species between females confronted to contrasted plant environments: pure maize, pure mugwort and mixture of maize and mugwort plants. We further related the differentially expressed (DE) genes to the host preference in ECB and ABB and investigated the functional categories involved. In parallel to this exhaustive *de novo* transcriptomic survey, and because of the importance of chemosensory genes which are considered as good candidates for the host recognition before oviposition, we conducted a specific analysis on a chemosensory genes subset previously characterized in ECB, ABB and other related *Ostrinia* species. As for *de novo* transcripts, we measured the gene expression differences of these chemosensory genes between ECB and ABB, and within each moth species, between contrasted plant environments. The discovery of candidate molecular mechanisms combined with our knowledge about the specialization patterns offers an opportunity to better understand the role of the host plants in the divergence between ABB and ECB.

## Material and methods

### Moths’ sampling and rearing

In the present study, we sequenced RNA from adult moths sampled from a previous experiment (Orsucci *et al*., 2016), where phenotypic and behavioral traits of moths facing different plant environments were measured. The whole experimental framework is reminded in Box 1 (“experimental design” session). Briefly, around 20 males and females were released in three types of conditions: (*i)* ‘choice’ cages, with maize and mugwort plants, (*ii)* ‘maize’ cages (usual host for ECB) and (*iii)* ‘mugwort’ cages (usual host for ABB). After 72h, several life history traits and particularly the egg laying were quantified. After measurements, all adults were recaptured and the females were used for the acquisition of expression data by RNA sequencing (see Fig. Box 1).

### Dissection, RNA extraction and sequencing

As we primarily targeted genes involved in the host preference and the host avoidance during oviposition, we focused on females only. Moreover, we assumed that these traits certainly involved chemosensory genes. Chemical reception is mediated by specialized sensory neurons located in antennae, mouthparts or legs, but chemosensory genes are also involved in non-sensory functions and expressed in other tissues (Wu *et al*., 2016). Thus, we chose to split head-thorax (HT) and abdomen (ABDO) tissues. HT is enriched in legs, antennae, and mouthparts while ABDO is enriched in digestive and genital organs. In addition, separate pools permitted increasing the sequencing coverage per pool of tissue. In total, we have dissected 220 females on ice to prevent thawing and RNA degradations. Total RNA was extracted from each tissue and individual with the AllPrep DNA/RNA 96 Kit (Qiagen), according to the manufacturer’s protocol. RNA quality and quantity were assessed using a Nanodrop, and equimolar quantities of individual extracts were pooled to obtain 24 samples corresponding to 2 repetitions x 3 experimental conditions x 2 tissues x 2 moth species. Each pool contained from 7 to 11 females depending on the available RNA samples (Table S1). Twenty-four cDNA libraries were constructed with the TruSeq Stranded mRNA Sample Preparation Kit (Illumina) following manufacturer’s protocol and sent to MGX platform (Montpellier, France) for single end 1 x 50bp sequencing on 6 lanes of a HiSeq 2000 (Illumina).

### *De novo* transcriptome assembly and annotation

After sequencing, all reads were subjected to quality control and trimming using Trimmomatic v0.25 to remove Illumina sequencing adapters and low quality reads (Bolger *et al*., 2014). Ribosomal and bacterial contaminants were removed using sortmeRNA v2.0, and Bowtie2 (Langmead & Salzberg 2012; version 2.2.4) mapping against a bacteria subset dataset extracted from NCBI database (October 2014).

To obtain two comprehensive transcriptomes from ECB and ABB libraries, we first assembled the high quality reads of the 12 ECB libraries on one side and of the 12 ABB libraries on another side using the trinity program suite (Haas *et al*., 2014) using default parameters and normalization. Then, for each of the resulting ECB and ABB assemblies, we included 454 data previously obtained by Gschloessl *et al*. (2013) from 4 ECB and 4 ABB adults, respectively. To achieve this goal, we used the EMBOSS (Rice et al 2000) splitter program (version 6.6.0) to create short overlapping subsequences (length 300 bp, overlap 200 bp) for each of the 454 transcripts and the HiSeq transcript sets, respectively. All 454 and HiSeq subsequences were then assembled with Newbler (454 Life Sciences Corporation, v 2.9) with 40 bp minimum overlap length, 97% minimum overlap identity and the ‘Het’ (for heterozygote) options. Finally, we selected the longest transcript as representative sequence for each gene to circumvent the occurrence of putative alternative transcripts that might bias expression analyses.

For functional and GO-terms annotations, a custom script was developed to compare ECB-ref and ABB-ref genes to NCBI nr (version may 2016) and UniprotKb (Bateman *et al*. 2015, version June 2016) with *blastx* BLAST (Altschul *et al*., 1990) and InterProScan (Jones *et al*., 2014).

### Candidate sensory genes

In parallel to the *de novo* transcripts’ reconstruction, we compiled a set of 141 reference genes involved in sensory perception previously described in ECB, ABB and related species of the *Ostrinia* genus (Leary *et al*., 2012; Miura *et al*., 2009; Miura *et al*., 2010; Wanner *et al*., 2010; Yang *et al*., 2015) composed of: odorant receptors (ORs), gustatory receptors (GRs), ionotropic receptors (IRs) sensory neuron membrane proteins and odorant degrading enzymes. As some of them may be orthologs between species, or similar between the studies, we removed the redundant sequences as identified by a reciprocal blast analyses. After this filtering step, the sensory dataset was reduced to 139 unique sequences (Table S2). This gene set, hereafter called ‘Senso-ref’, was used as an independent gene reference from ABB-ref and ECB-ref to focus on sensory genes.

### Raw reads mapping and normalization

Raw reads were mapped onto ABB-ref, ECB-ref and Senso-ref using Bowtie2 (Langmead & Salzberg 2012; version 2.2.4), with default parameters and “sensitive” option. After the mapping, we produced the counts of mapped reads per gene with samtools program (Li *et al*., 2009). Transcripts-specific variations (dispersion and logarithmic fold-changes of reads counts among experimental conditions) were estimated with the Bioconductor edgeR (Empirical Analysis of Digital Gene Expression Data in R) package (Robinson *et al*. 2010; version 3.8.6). A negative binomial distribution was chosen as model for counts variations, and transcripts with less than one count per million for at least half of libraries were removed. Reads counts were normalized among conditions (*i.e*. among sequenced libraries) with the EdgeR function *calcNormFactors* and TMM option (Robinson and Oshlack, 2010) and log (base 2) fold changes (Log2FC) were calculated for each gene between conditions.

In addition, a principal component analysis (PCA) was conducted on the normalized read counts as implemented in the FactoMineR library of R (Lê *et al*., 2008). The following categorical variables were input as supplementary variables: the moth species (ABB *vs*. ECB), the experimental condition (choice, mugwort, maize), and the tissue (ABDO *vs*. HT).

### Identification of differentially expressed (DE) genes: response to environment and divergence between ECB and ABB

The main source of variation in expression was, as expected, explained by the HT *vs*. ABDO tissues rather than by species or experimental conditions (Fig.1). Thus, we analyzed ABDO and HT libraries separately. The variations of read counts by transcript (*t_i_*) were analyzed with linear model as implemented in EdgeR, to estimate the statistical support of the experimental effects. We combined the two explanatory variables, *moth species* and *experimental condition*, into a so-called ‘*modalities’* factor, and applied the following model:

> Model 1: *t_j_* = *modalities*.

**Fig. 1.**
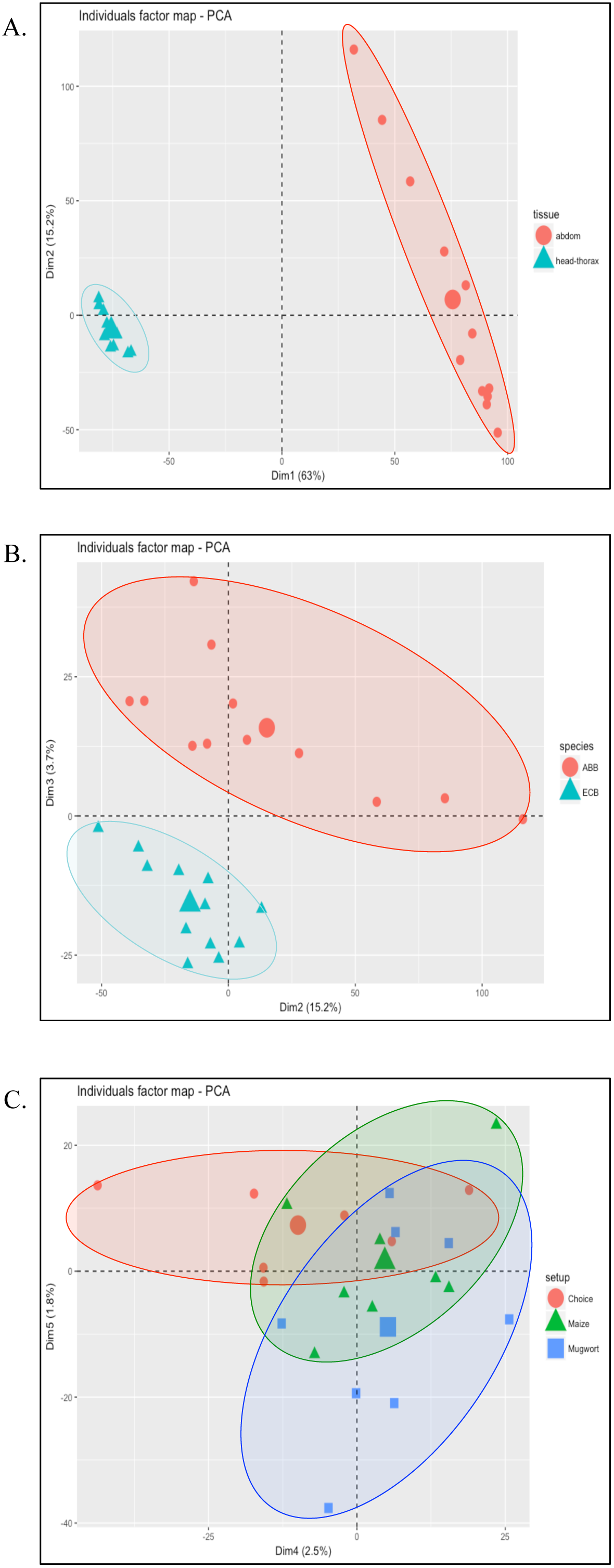
PCA individual factor map of mean raw counts of 9413 genes in 24 RNA libraries. A: plot on 1-2 axes, ellipses circumvent libraries mean raw counts per tissue (ABDO *vs*. HT); B: plot on 2-3 axes, ellipses circumvent libraries mean raw counts per moth species (ECB *vs*. ABB); C: plot on 4-5 axes, ellipses circumvent libraries mean raw counts per experimental condition (choice *vs*. maize *vs*. mugwort). The percentage of explained variation is indicated on each dimension axis.

We then used a contrast analysis to identify *i)* transcripts whose expression varied significantly between the ABB and ECB whatever the experimental condition, and *ii)* transcripts whose expression varied significantly between experimental conditions (three comparisons: maize and mugwort; maize and choice; mugwort and choice) whatever the moth species. Transcripts were considered differentially expressed (DE) in the whole experiment if they had a false discovery rate (FDR) adjusted *p-value* < 0.01.

Gene ontology (GO) enrichment was calculated in R using “*top GO*” package (v2.26.0) from the Bioconductor project (Alexa and Rahnenfuhrer, 2010). We used the GO annotation described previously as ‘custom input’. We removed GO terms with fewer than 5 annotated genes (using the *nodeSize* parameter set to 5). We used the ‘weight’ algorithm and the Fisher exact test procedure. We report enriched GO categories with *p-values* < 0.01 for the following DE gene lists: DE genes between ECB and ABB species, and DE genes between experimental environment (maize and mugwort, maize and choice, mugwort and choice).

### Categorization of differentially expressed (DE) genes according to behavioral traits

We further refined the categorization of DE genes to relate them to the preference and avoidance traits and behaviors previously observed in Orsucci *et al*. (2016) (Box 1). For that, we split the data set and analyzed separately ECB and ABB data through the simplified model:

> Model 2: *t_i_* = *experimental condition*

and we compared *i)* gene expression between mugwort samples and the mean expression of the two other experimental conditions in ECB sub-data, to detect genes potentially involved in maize preference during oviposition in ECB (hereafter referred as ECB-pref); *ii)* gene expression between maize samples and the mean expression of the two other experimental conditions in ABB sub-data to detect genes involved in mugwort preference during oviposition in ABB (ABB-pref); and *iii)* gene expression between maize-only and the mean expression of the two other experimental conditions in ECB sub-data to detect the genes involved in mugwort avoidance during oviposition in mugwort environment in ECB (ECB-avoid).

For all models, the *p-values* were adjusted for multiple testing using the Benjamini-Hochberg procedure (FDR < 0.01). Gene ontology (GO) enrichment was calculated as previously, on ECB-pref, ABB-pref and ECB-avoid gene lists.

## Results

### ECB- and ABB-ref assemblies and annotation

The main features of the raw read data and ECB and ABB assemblies (HiSeq only or HiSeq-454 combined) are detailed in Table 1. We observed that the combination of published 454 transcripts and *de novo* HiSeq transcripts has doubled the length of transcripts (Table 1) compared to the solely *de novo* HiSeq assembly. Taking the longest sequence as unique represent for each unigene resulted in 9,415 ECB genes and 9,992 ABB genes (Table 1). Homology searches in genomic and protein databases yielded significant hits for 61% of the ECB genes among which 80% had an associated GO term. More than 97% of the annotated genes were homologous to Lepidoptera genes, followed by homologs to other insect orders (Table S3). To a much lower extent, hits with genes of plants, viruses and bacteria were also observed. Among microorganism, viruses were the most prevalent. Details about their detection and annotations are given in Table S4A.

**Table 1.** Sequencing data and principal features of ECB and ABB assemblies

|  |  |  | **ECB** | **ABB** |
| --- | --- | --- | --- | --- |
| Number of Raw reads | HiSeq | after sequencing | 632,618,343 | 655,684,618 |
|  |  | after filtering | 466,145,250 | 498,030,304 |
| Number of unigenes | HiSeq |  | 41,155 | 42,913 |
| Number of transcripts | HiSeq |  | 41,212 | 42,969 |
|  | HiSeq-454 | all transcripts | 11,456 | 12,024 |
|  | HiSeq-454 | longest transcript per unigene | 9,415 | 9,992 |
| Transcript N50 length (bp) | HiSeq |  | 530.41 | 544.16 |
|  | HiSeq-454 |  | 1,221 | 1,260 |
| Assembled transcriptome (Mbp) | HiSeq-454 |  | 13.7 | 15.2 |

### DE genes in ECB- and ABB-ref: global pattern

The DE genes detection was done in parallel on ECB-ref and ABB-ref. Quantitative and qualitative trends were highly correlated between both analyses (data not shown), so that, for simplicity, we present in the following the results only for ECB-ref. We chose to conduct two analyses rather than constructing a mixed reference by pooling ECB and ABB reads, in order to avoid chimeric transcripts. At the global analytic scale, we did not observe noticeable differences in the number or nature (function and GO term) of the DE genes. Nonetheless at a finer scale (*e.g*. for one given gene), the ECB and ABB reconstructions may differ.

The overall percentage of mapped reads on the ECB-ref transcripts varied from 64 % to 71 % among the 12 HT RNA libraries and from 58% to 65% among the 12 ABDO libraries (Table S1). The proportion of multiple mapped reads was less than 1% in HT (0.51%) and ABDO (0.39%) libraries. After filtering genes with coverage lower than or equal to 1X, we identified a total of 9326 and 9186 genes expressed over all experimental conditions in the HT and ABDO samples, respectively. Among them, a majority (around 98% in HT and 95% in ABDO samples) was shared among all experimental conditions (Fig. S1). We detected 24 and 17 private genes in ECB samples and 19 and 118 private genes in ABB for HT and ABDO samples, respectively. After Fisher’s exact test, we did not found significant enrichment in GO categories in the private genes sets.

### DE genes and source of variation: species and experimental effects

Set apart from the tissues, the main source of differential expression was due to contrast between ECB and ABB samples (Fig. 1B). In HT samples, we detected 337 and 268 transcripts significantly over- and under-expressed in ECB *vs*. ABB. For ABDO samples, 828 and 132 transcripts were significantly over- and under-expressed in ECB *vs*. ABB (Table 2, model 2).

**Table 2.**
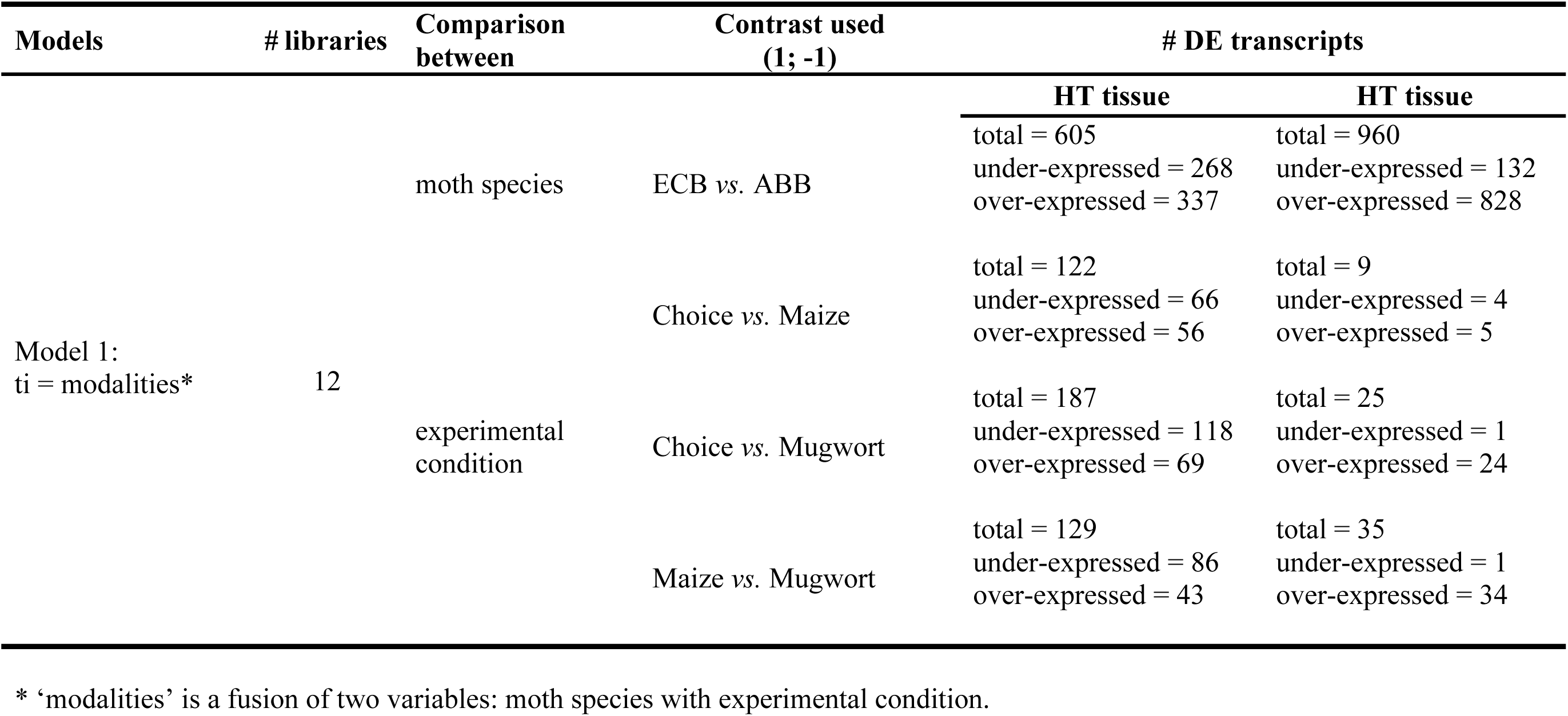
GLM model 1.

Annotated DE genes between ECB and ABB (Table S5), and expressed in HT tissues, were mostly enriched in GO terms (Table S6) relating to: *i)* DNA/RNA metabolic processes (*e.g*. reverse transcriptases, transposases, pol proteins and transferases) and *ii)* organic compounds metabolic processes (*e.g*. fructofuranosidases and UDP-glucuronosyltransferase). In the most DE genes (|log2FC| > 2) we also observed several cytochrome P450s involved in detoxification processes, one of which was previously characterized in *O. furnacalis* (gb|ACF17813.2) to the DIMBOA toxic compound of maize, and another was an immune-induced protein previously characterized in *Ostrinia* species (gb|AGV28583.1, Table S5). In ABDO tissues we observed enrichment in GO terms relating to DNA metabolic processes, molecular binding and transport (Table S6). In the most DE genes (|log2FC| > 5, Table S5) we identified apolipophorins potentially involved in immune response (Zdybicka-Barabas & Cytryñska 2013) as well as the DIMBOA-induced cytochrome P450 previously described in *Ostrinia furnacalis* (gb: ACF17813.2, |log2FC| > 5, Zhang *et al*. 2016).

The experimental condition explained a smallest part of the variation in gene expression (Table 2, model 2, Fig. 1C). From 122 to 187 genes were differentially expressed between experimental condition comparisons (choice *vs*. mugwort, choice *vs*. maize and mugwort *vs*. maize) in HT samples. Such DE genes were much fewer in ABDO samples (9 to 35). These contrasted number of DE genes reflected the higher between-repetition dispersion and between-experimental condition overlap in ECB ABDO tissues compared to ECB HT samples (Fig. 2A-B), and also in ABB samples *vs*. ECB sample (Fig. 2C-D *vs*. Fig. 2A-B).

**Fig. 2.**
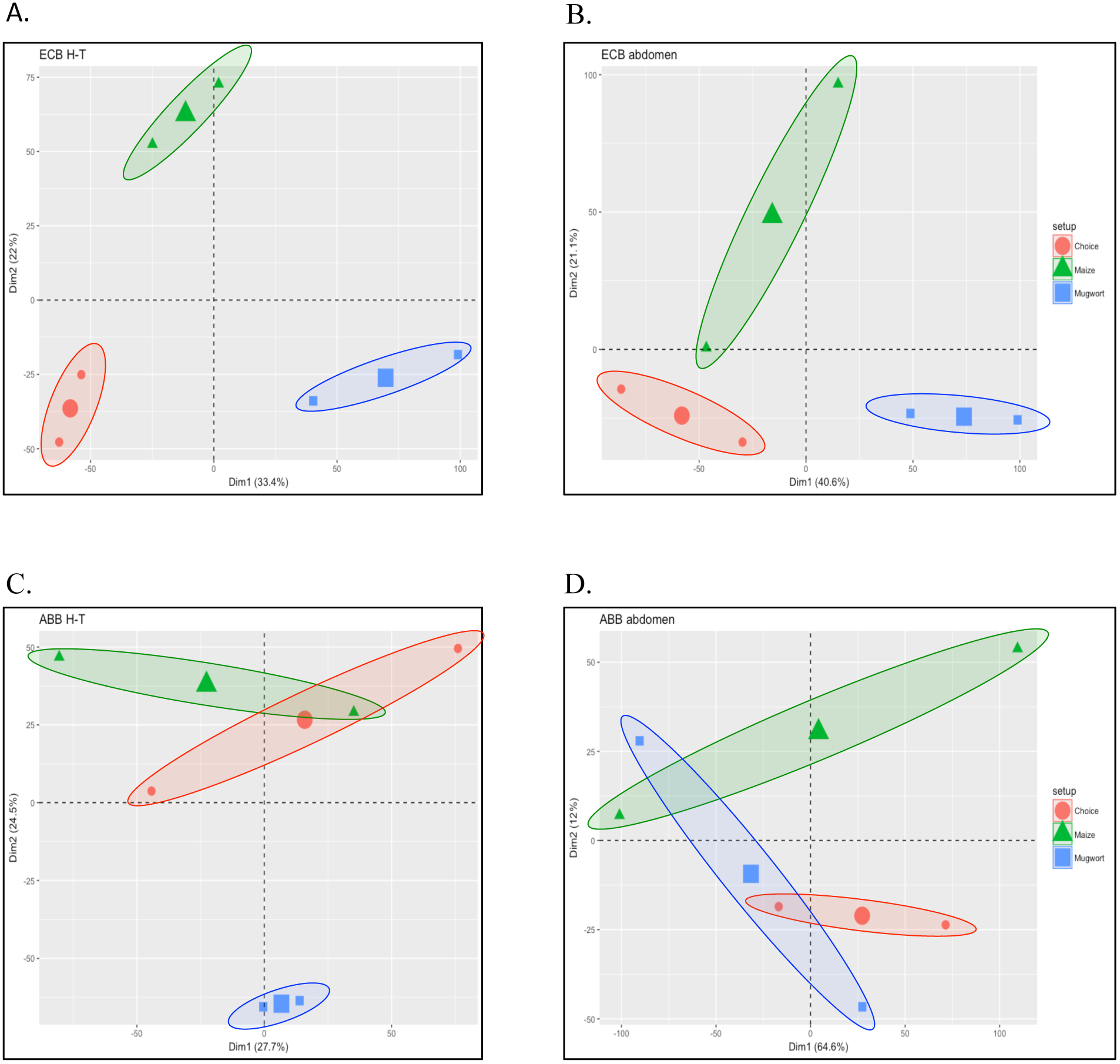
PCA individual factor map of mean raw counts of 9413 genes in 6 RNA libraries. A: ECB HT samples; B: ECB ABDO samples, C: ABB HT samples; D: ABB ABDO samples. The percentage of explained variation is indicated on each dimension axis.

Annotated DE genes between experimental conditions (Table S7) were enriched in a wide variety of biological process GO categories (Table S6) covering feeding and mating behavior, sensing, molecule transport and binding, organic compounds and DNA/RNA metabolism. In the most DE genes (|log2FC| > 5, Table S7) of HT and ABDO samples, we identified various enzymes potentially involved in digestion, detoxification and immunity (*e.g*. glucose dehydrogenase, valine-rich midgut proteins [VMP], heat shock proteins, and cytochrome P450) and one protein potentially involved in sensing (transient receptor potential protein).

### DE genes and behavioral traits

We then focused on DE genes potentially related to specific traits and behaviors observed at the phenotypic level (Box 1), by a specific contrast analysis in the GLM procedure (Table 3). With this analysis, we categorized the DE genes identified by the Model 3 (Table 3) in maize preference in ECB (ECB-pref), mugwort avoidance in ECB (ECB-avoid) and mugwort preference in ABB (ABB-pref). Remarkably, much more DE genes were observed in ECB ([260-786] for ECB-pref and [132,149] for ECB-avoid; Table 3 and S8) than in ABB ([4-56] for ABB-pref, Table 3 and S8). Moreover, the preference genes are in majority private to ECB or ABB (Fig. S1).

**Table 3.** GLM model 2.

| Targeted trait | # libraries | Models | Contrast used<br>(0.5; 0.5; -1) | # DE transcripts |  |
| --- | --- | --- | --- | --- | --- |
|  |  |  |  | HT tissue | ABDO tissue |
| Maize preference in ECB | 6 | Model 3:<br>ti = experimental<br>condition | Choice and Maize<br>vs.<br>Mugwort | total = 260<br>under-expressed = 148<br>over-expressed = 118 | total = 786<br>under-expressed = 608<br>over-expressed = 178 |
| Mugwort avoidance in<br>ECB |  |  | Choice and Mugwort<br>vs.<br>Maize | total = 132<br>under-expressed = 84<br>over-expressed = 48 | total = 149<br>under-expressed = 128<br>over-expressed = 21 |
| Mugwort preference in<br>ABB | 6 |  | Choice and Mugwort<br>vs.<br>Maize | total = 56<br>under-expressed = 15<br>over-expressed = 41 | total = 4<br>under-expressed = 3<br>over-expressed = 1 |

We observed enrichment in various GO categories for the DE ECB-pref, ABB-pref and ECB-avoid genes, related to the metabolism of carbohydrates, lipids, proteins, and amino acids, the detoxification mechanisms, the immunity and the chemosensory repertoire (Table S6). In these DE gene lists, we last considered those for which the expression also differed between ECB and ABB. They include a reverse transcriptase, various enzymes involved in digestion and detoxification (*e.g*. mucin, Dzip1, Beta-fructofuranosidase, phospholipase, serine protease), one immunity protein (lysozyme precursor), a heat shock protein and one egg yolk protein (Table S5). All these DE genes were involved in the ECB-pref and ECB-avoid, making them the best candidates to the divergence driven by the specialization to the maize.

### DE transcripts over sensory candidate genes

One hundred and nine chemosensory genes over the 139 Senso-ref candidates were expressed in at least one library (Table S9). As for the *de novo* transcriptome, most of the variation in expression for these genes was due to the tissue type (ABDO *vs*. HT) with 65 DE genes, followed by the moth species (ECB *vs*. ABB) with 36 DE genes. Globally, 28 genes were differentially expressed according to the experimental conditions, among which a majority (24) differed between pure mugwort and pure maize conditions. Focusing on DE genes candidate potentially involved in traits and behaviors (ABB-pref, ECB-pref and ECB-avoid) observed at the phenotypic level (Box 1, Table S9), by the specific contrasts analysis in the GLM procedure, we highlighted eight genes involved in maize preference in ECB samples: four odorant receptors (ORs), one aldehyde oxidase, one carboxylesterase, and one ionotropic receptor (IR). Only two genes (ORs) were involved in mugwort avoidance in ECB samples, and three genes (ORs) in mugwort preference in ABB samples. Notably, genes involved in ECB preference differed from those involved in ABB preference and in ECB avoidance (Table S8). Finally, among the 28 genes differentially expressed according to the plant environment, 11 (five ORs, five IRs and one caroboxylesterase) were also differentially expressed between ABB and ECB samples.

## Discussion

By using a combination of a RNAseq approach without *a priori* and a specific analysis targeting chemosensory genes, we searched for genes expressed in ECB and ABB females that were exposed to various environment with or without their usual host plant. We identified genes and functions differentiating ECB and ABB whatever, or depending on, the environment. Moreover, we classified these DE genes according to their expression pattern to relate them to preference and avoidance behaviors in ECB and ABB gravid females. We hence identified candidate genes to species divergence by host specialization.

### Response to plant environment in ECB and ABB during oviposition: global pattern

In average over all transcripts, we first observed that more genes were up or down regulated in HT than in ABDO tissues. This observation corroborated the expectation that sensory and neuronal signals, which are active mainly in HT organs, are major factors in oviposition (Reisenman and Riffell, 2015; Trona *et al*., 2013). Second, we recorded more DE genes in ECB than in ABB samples. We may assume that this result is a signature of the strength and variability of the host specialization in ECB (combining host preference and alternative host avoidance), while ABB appeared less specialized to its own host in terms of physiological and behavioral traits (Orsucci *et al*., 2016 and Box 1). Last, we found that the genes involved in the preference for their respective host were different between ECB and ABB, as well as those involved in mugwort rejection in ECB, suggesting that the ECB shift onto maize was characterized by genetic novelties.

At the functional level, the response to plant environment in ECB and ABB during oviposition involved many processes, including metabolism of carbohydrates, lipids, proteins, and amino acids, detoxification mechanisms, immunity and chemosensory repertoire. These functions are key processes of the host specialization in phytophagous insects (Simon *et al*., 2015), but were until now mainly illustrated at the larval stage (Eyres *et al*., 2016; Hoang *et al*., 2015; Koenig *et al*., 2015; Li *et al*., 2013; Ragland *et al*., 2015; Silva-Brandão *et al*., 2017). We provide in the present study novel results for the adult stage, revealing some other candidates of the host specialization like takeout-like proteins, previously reported as involved in circadian rhythms and feeding behavior in *Drosophila* (Sarov-Blat *et al*. 2000).

### The chemosensory genes in host choice: new candidate genes and divergence between ECB and ABB

The analyses targeted on chemosensory genes supported their implication in host discrimination in ECB, ABB or both. Moreover five ORs, five IRs and one caroboxylesterase appeared as suitable candidates for their putative role in species divergence by specialization. Indeed, their expression responded to plant environments and, at the same time, differed between moth species. Among them, one OR (LC002733.1, Table S9) was particularly relevant, because it clearly discriminated maize and mugwort conditions in ECB, either for the maize preference or for the mugwort avoidance. In contrast with lot of other OR involved in pheromone recognition by males (Wanner *et al*., 2010), Yang *et al*. (2015) have observed that this OR was higher expressed in the female in ECB. We highlight that the DE open the way toward new candidates potentially involved in plant odor discrimination in females during oviposition. Last, we identified among the genes differentially expressed between moth species and which are involved in the ECB-pref, one OR (JN169130, Leary *et al*. 2012) previously reported as more expressed in males than females in ECB and ACB (Asian corn borer, another sibling species of *Ostrina* complex species). This receptor may thus be an example of pleiotropy or synergetic activity for the pheromone recognition in male and plant recognition in females. An essential requirement to speciation via specialization is the existence of a link between reproductive isolation and specialization due to divergent selection for different environments (Matsubayashi *et al*., 2010b). In ECB and ABB, two pheromone lineages coexist and are partially reproductively isolated due to the discrimination by males of the pheromone blends emitted by females (Glover *et al*., 1987; Lassance, 2010). A gene implicated simultaneously in the pheromone recognition in males and host plant discrimination permitting the oviposition choice by the females would be a direct link for mating isolation and host specificity (Via and Hawthorne, 2002).

### A major role of metabolism of organic compounds during oviposition and in divergence between ECB and ABB

Beside recognition of plant chemical cues, the analysis based on whole of *de novo* assembly highlighted a huge variety of other functional categories involved either moth species divergence, or in host plant choice or in both.

The organic metabolic processes represented by key genes previously identified and involved in the differentiation between ECB and ABB (Orsucci *et al*. submitted) but also in various phytophagous lepidopterans (Simon *et al*., 2015) highlighted the importance of the digestion and detoxification of plant metabolites in particular at the larval stage. At the adult stage, chemosensory processes have been emphasized, because ovipositing females primarily use plant secondary metabolites as cues to locate and select suitable plants (Städler, 2008). Our study suggests that digestion and detoxification of non-volatiles compounds are additionally important factors in the host choice. This result may be unexpected because ECB is often considered as a ‘capital breeder’, *i.e*. as an insect that feed only during the larval stage, as it is often the case for lepidopterans (Bonoan *et al*., 2015). Our results and previous works (Derridj *et al*., 1992; Leahy and Andow, 1994) show, on the contrary, that nutrition (and in particular carbohydrates) may play a key role in the oviposition behavior, egg weight, fecundity and female longevity.

Interestingly, the activation of genes belonging to specific metabolic pathways differed between ECB and ABB, with for examples: mucin, serine protease and phospholipase genes involved in ECB-pref and ABB-pref. These genes hence require more investigation to test if they reflect differences in metabolic strategies employed by ECB and ABB to survive, develop, and reproduce on their respective host.

### The unexpected actors: virus and retrovirus-like transposon

Besides the metabolism of organic compounds, we also observed in the most DE genes, functions relating to detoxification (cytochrome P450) and immune-induced protein previously characterized in *Ostrinia* species (gb|AGV28583.1, Table S5). These immune and detoxification genes may be mobilized to detoxify some plant metabolites but also involved in the defenses against bacteria and virus. Indeed, we observed the occurrence of plants, viruses and bacteria genes, which are probably due to some rests of the food bolus or the presence of environmental, commensal, symbiotic, or pathogenic microorganisms. The RNA sequencing was not designed to target the microorganism community, so that a fine and quantitative analysis would be unwarranted. Yet the presence of some of this alien fauna impacted the gene expression of ECB and ABB females, as illustrated for example by the activity of DNA-directed RNA polymerases (Table S5), implicated in the replication of RNA virus. In particular for virus, we observed contrasted patterns between ECB and ABB or between experimental conditions, with a higher prevalence of cypovirus-like genes in ABB head-thorax samples in maize conditions than in ECB and other conditions (Table S4A). We thus suggest that immune and defense genes differentially expressed between ECB and ABB, or between maize/mugwort conditions, may have been activated to deal against pathogens present in the experimental environment. Interestingly, the prevalence of these microorganisms as well as the defense response of female moths differed between ECB and ABB samples, and between environments. These results, and previous similar observations at the larval stage (Orsucci *et al*. submitted), warrant further studies on other moth populations and plant species, to *i)* evaluate the variability and structuration of the microorganism communities *in natura, ii)* measure the immune response and its divergence between ECB and ABB, and *iii)* characterize the interactions between host plant, microorganisms and species divergence. Such pluritrophic interactions have been demonstrated in different studies (see Raymond *et al*., 2002). They deserve further investigations in the specific context of the ECB-ABB divergence.

In parallel, we also observed the expression of retroviral like transposons (Table S4B) and of the various genes associated to their transposition and spread, in particular in HT tissues of ECB. The putative role of transposable elements (TE) in ECB and ABB divergence has not yet been reported, but these elements are known to interfere with adaptive response to various environmental stresses in others and various taxa (Casacuberta and González, 2013). The TE burst may thus in some circumstances be a driver of speciation events (Belyayev, 2014). In ECB, Lillehoj (1991) proposed a scenario in which the aflatoxin (toxin produced by the microbial inhabitant of the insect digestive tract) induced the activation of mobile genetic elements such as transposons, plasmids or chromosomal mutation, in response to environmental changes. According to this author, aflatoxin-producing fungi may be symbiotic in certain equilibrium conditions and pathogenic in conditions of nutrient imbalance, notably in presence of elevated levels of digestible carbon compounds. The ecological disequilibrium induced by crop monoculture is supposed to have favored ECB populations chaining two or more life cycles per crop rotation over univoltine populations prone to a diapause period. According to Lillehoj (1991), the aflatoxin and its properties on host DNA and transposons activation may have facilitated this novelty in ECB. No similar mechanism has been described in ABB, and further studies are required to test directly the role of the transposons activity on ECB-ABB divergence and on maize colonization by ECB.

## CONCLUSION

Our results allowed identifying genes and functions candidates for specialization and for species divergence between ECB and ABB. By an ad-hoc categorization, we discriminated between genes responding to the environment with similar or divergent pattern in ECB and ABB. Independent approaches such as functional or population genomics will be required to confirm their role in the history of ECB host shift onto maize. In this perspective, we confirmed the role of sensing processes and identified new candidates in the chemosensory repertoire. These candidates should be tested in different populations and contexts, since they could help designing efficient control tools for the crop pest management (Reisenman *et al*., 2016). Finally, we also highlighted other lines of research on carbohydrates metabolism or virus and retrovirus dynamics, which are also promising ways to better understand the divergence process between cryptic sibling species.

### Availability of data

The data (raw RNAseq read and the assembled ABB and ECB transcriptomes) have been deposited with links to BioProject accession number PRJNA396660 in the NCBI BioProject database (https://www.ncbi.nlm.nih.gov/bioproiect/).

## Acknowledgments

We thank Béatrice Ramora, Anne Zanetto and Christophe Reynaud for technical assistance in plant cultivation in DIASCOPE (experimental research INRA station) site. We thank also Denis Bourguet, Pascal Milesi and Carole Smadja for their critical and helpful comments on the manuscript. This work received supports from ANR Adapt-Ome (ANR-13-BSV7-0012-01) to RS and DB, from the INRA department ‘Santé des Plantes et Environnement’ and from the University of Montpellier for MO PhD studentship.

## SUPPLEMENTARY TABLES AND FIGURES

The supplementary tables and figures are provided as separate files. The tables are provided as xlsx files named “Table S#.xlsx” and the figures as pdf files named “Fig. S #.pdf’.

**Table S1.** Sequencing characteristics of the RNA pools. Number of samples per pool, type of tissue (A: abdomen, HT: head-thorax), line ID of the Hiseq sequencing, number of raw reads, and percentages of read mapping on ECB-ref and ABB-ref.

**Table S2.** Published chemosensory *Ostrinia* spp. genes. This gene-set is used as reference for targeted read mapping. Issued from Yang *et al*. (2015), Leary *et al*. (2012), Wanner *et al*. (2010), Miura *et al*. (2009) and Yasukochi *et al*. (2011)

**Table S3.** Top hit species after BLAST analysis. For the ECB-ref transcripts (sheet 1) and ABB-ref transcripts (sheet 2)

**Table S4.** Detailed information for DE genes with homology with virus (A) and retrovirus-like transposon sequences (B). Mean normalized read counts per experimental set-up, species and tissues; nr, Uniprot and GO annotations.

**Table S5.** Detailed information for genes DE between ECB and ABB species. Log2FC (only|Log2FC| > 2 are presented); nr, Uniprot and GO annotations; mean normalized read counts per experimental set-up, species and tissue. In blue: Log2FC < -2; in red: Log2FC > +2. For clarity, transcripts without Blast hits and transcripts with anonymous annotation (*e.g*. ‘uncharacterized protein’) were removed.

**Table S6.** Fisher test of enrichment in GO term categories between the set of DE transcripts (test set) and the whole transcriptome (reference set). Sheet 1: enrichment in genes DE between species; Sheet 2: enrichment between experimental environments (maize, choice, mugwort); Sheet 3: enrichment in ECB-pref, ECB-avoid and ABB-pref gene lists. The first column indicates the sample and tissue tested (in grey: ABDO, no colour: HT), and the BP (biological process) and MF (molecular function) category of the associated GO term.

**Table S7.** Detailed information for genes DE between experimental environments (maize, choice, mugwort). Log2FC (only|Log2FC| > 2 are presented); nr, Uniprot and GO annotations; mean normalized read counts per experimental set-up, species and tissue. In blue: Log2FC < -2; in red: Log2FC > +2. For clarity, transcripts without Blast hits and transcripts with anonymous annotation (*e.g*. ‘uncharacterized protein’) were removed.

**Table S8.** Detailed information for genes related to ECB-pref, ECB-avoid and ABB-pref. Log2FC (only|Log2FC| > 2 are presented); nr, Uniprot and GO annotations; mean normalized read counts per experimental set-up, species and tissue. In blue: Log2FC < -2; in red: Log2FC > +2. For clarity, transcripts without Blast hits and transcripts with anonymous annotation (*e.g*. ‘uncharacterized protein’) were removed.

**Table S9.** Detailed information for DE chemosensory genes. |Log2FC| > 2 are presented in ECB-pref, ABB-pref, ECB-avoid, experimental effect and species effect.

**Figure S1. Venn diagram for ECB-ref (A and B) and ABB-ref (C and D) transcripts.** The Venn diagram indicate the number of transcript shared between the different experimental conditions for (A and C) the H-T samples and (B and D) the ABDO samples. The ECB samples are represented by heat colors and ABB sample by cold colors.

